# Genome sequence of the blue flowering *Centaurea cyanus*

**DOI:** 10.1101/2025.01.31.635922

**Authors:** Chiara Marie Dassow, Ronja Friedhoff, Boas Pucker

## Abstract

The genome of the cornflower (Centaurea *cyanus*) was sequenced with long reads (ONT) to reveal the full sequences of genes in the flavonoid biosynthesis. Of particular interest are genes responsible for the striking blue pigmentation of the flower. Through basecalling, read correction and assembling the sequenced DNA strands, we were able to generate a genome sequence with 98.8% BUSCO completeness, N50 of 16 Mbp, and an accuracy of 72 reported by Merqury. HERRO-based correction of the R10 nanopore sequencing reads resulted in a substantially improved assembly compared to uncorrected reads. The gene prediction revealed structural genes of the flavonoid biosynthesis and associated MYB and bHLH transcription factors. A tandem duplication of F3’H was discovered. Additionally, a putative tandem duplication resulting in three copies of the anthocyanin biosynthesis activating MYB was observed.

## Introduction

Cornflower, *Centaurea cyanus*, is a member of the *Asteraceae* and widespread in gardens and parks as an ornamental plant (Zhang *et al*., 2023). This plant is best known for the striking blue color of its radial symmetric flower (**Fig. 1**). The rarity of blue flowers in plants might contribute to the popularity of cornflowers as ornamental plants. Pigmentation in plant flowers fulfills multiple roles, from attracting pollinators to protecting themselves against oxidative stress (Heim *et al*., 2002; Shen *et al*., 2022; Kellenberger & Glover, 2023; Grünig *et al*., 2024). While betalains and carotenoids play a role in petal colour formation, the most prominent pigments are flavonoids, specifically anthocyanins that confer colours to most flowers (Winkel-Shirley, 2001; Tanaka *et al*., 2008; Lu *et al*., 2024). Among them, only a specific group of anthocyanins can provide blue coloration. Delphinidin derivatives, characterized by a triple hydroxylation of the B-ring, are responsible for bluish color (Liu *et al*., 2022). Alternatively, structural colour or different forms of co-pigmentation can lead to a blue appearance (Glover & Whitney, 2010; Trouillas *et al*., 2016; Airoldi *et al*., 2019). One form of co-pigmentation is the formation of metal ion complexes as it is seen in *Salvia patens* or *Meconopsis grandis* (Takeda *et al*., 1994; Yoshida *et al*., 2006). The formation of colourful metal complexes is rare in living organisms; only a few species have been described to carry a metal ion complex, turning them blue (Takeda, 2006). One famous example is the protocyanin in the cornflower (Takeda *et al*., 2005). This complex comprises 6 cyanin molecules (anthocyanin), 6 apigenin molecules (flavone), 2 calcium ions, one magnesium ion, and one ferric ion (Shiono *et al*., 2005).

**Fig. 1:**
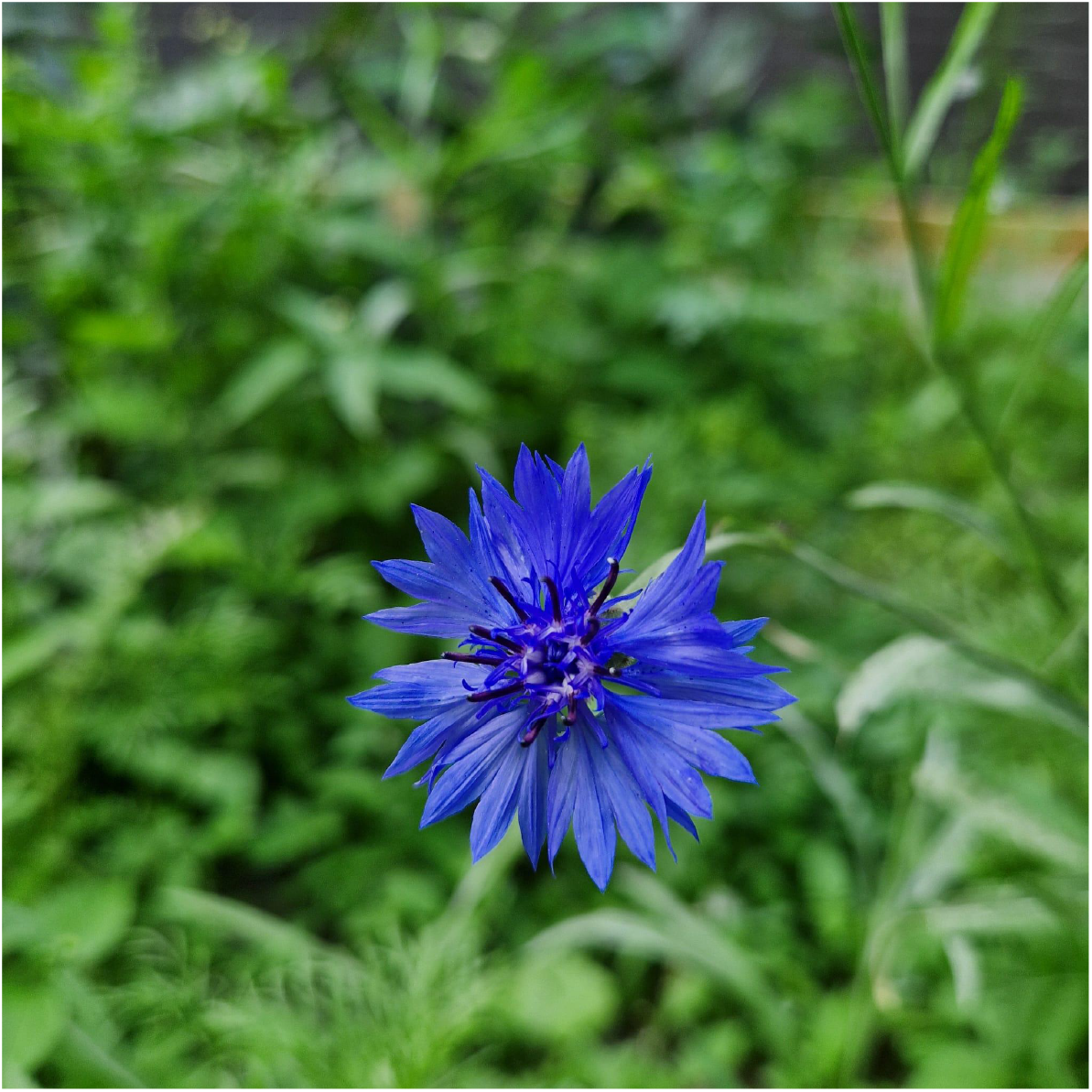
Striking blue flower of a *Centaurea cyanus* plant.

Anthocyanins and flavones are produced by two distinct branches of the flavonoid biosynthesis (Winkel-Shirley, 2001). Flavonoids are generally based on phenylalanine which is channeled through the general phenylpropanoid pathway. The chalcone synthase (CHS) catalyzes the first committed step of the flavonoid biosynthesis (Ferrer *et al*., 1999). Chalcone isomerase (CHI) and flavanone hydroxylase (F3H) are catalyzing the next steps in the flavonoid biosynthesis (Forkmann *et al*., 1980; Jez *et al*., 2000). The flavone biosynthesis is branching off with flavone synthase II (FNS II) being the committed step in most plant species (Martens & Forkmann, 1999). Only the Apiaceae harbour an alternative gene namely flavone synthase I (FNS I) which encodes an independently evolved flavone synthase (Martens *et al*., 2003; Pucker & Iorizzo, 2023). Dihydroflavonol 4-reductase (DFR), anthocyanidin synthase (ANS), anthocyanin-related glutathione S-transferase (arGST), and an UDP-dependent glycosyltransferase (UFGT) are involved in the anthocyanin formation (Davies *et al*., 2003; Turnbull *et al*., 2003; Yonekura-Sakakibara *et al*., 2012; Eichenberger *et al*., 2023; Grünig *et al*., 2024).

Recent studies have shown the existence of anthocyanin regulating MYBs and bHLHs in the cornflower (Deng *et al*., 2020). These are transcription factors needed, among other factors for the regulation of flavonoid biosynthesis genes (Xu *et al*., 2015). First analysis of flavonoid biosynthesis genes have been done, but the sequences of respective genes have not been published (Deng *et al*., 2019).

The cornflower flower itself has been reported to have many health benefits, as it increases the uptake of protein, is high in antioxidants as well as in fibers and can function as an all natural dye (Król *et al*., 2024). Harvesting these properties for our day to day life could be advantageous to humans, but also be more sustainable for nature. Elucidating the responsible mechanism for the high antioxidative character of protocyanin provides a solid ground for further developments from introducing colour into other pigmented plants to usage in our food.

By sequencing the cornflower genome with long read sequencing technology, we were able to generate a highly contiguous genome sequence. Structural genes and transcriptional regulators required for the flavonoid biosynthetic pathway in the cornflower were identified. By finding these genes we are one step closer to generating flavonoids and flower colour synthetically. This study gives an insight into the genome of the cornflower and the relevant genes needed for the flavonoid biosynthesis leading to the blue flower colour.

## Materials & Methods

### Plant Sample and DNA Extraction

The high molecular weight DNA extraction from young leaves was performed based on a previously optimized CTAB-based protocol followed by established quality control steps (Siadjeu *et al*., 2020; Horz *et al*., 2024). In brief, DNA quantification via NanoDrop, DNA quality assessment on an agarose gel, and DNA quantification via Qubit were performed. Suitable samples were subjected to a short fragment depletion step using the Short Read Eliminator kit (PacBio).

### Nanopore sequencing of high molecular weight DNA

The library preparation for nanopore sequencing was conducted with 1 µg of DNA following the supplier’s instructions for SQK-LSK114. Sequencing was performed on R10.4.1 flow cells on MinION Mk-1B as previously described (Horz *et al*., 2024). Sequencing results were collected in pod5 files and transferred to a cloud environment for basecalling with GPU support. Different basecalling approaches were evaluated with respect to the output quality. Initially, basecalling was performed with Guppy v6.5.7 (ONT) in the high accuracy mode (dna_r10.4.1_e8.2_400bps_5khz_hac.cfg). An additional run was performed in the super high accuracy mode (dna_r10.4.1_e8.2_400bps_5khz_sup.cfg).

Basecalling with dorado v0.8.3 using the model dna_r10.4.1_e8.2_400bps_sup@v5.0.0 and detection of modified bases (--modified-bases “5mCG_5hmCG”) was performed for comparison. An additional correction of the reads with HERRO v1 (Stanojević *et al*., 2024) was performed.

### Genome sequence assembly

Genome sequence assemblies were generated with different tools to identify the best assembly strategy. A Shasta v0.13 (Shafin *et al*., 2020) assembly attempt with default parameters and based on the guppy basecalled data generated a very small genome sequence indicating an artifact. Shasta assembly attempts with the same settings were performed based on dorado-basecalled and HERRO-corrected data. NextDenovo v2.5.2 (Hu *et al*., 2024) was run with the following arguments based on the guppy-basecalled and dorado-basecalled data: genome_size=700m, seed_cutoff=10000, correction_options=-p 1 -max_lq_length 10000 -r ont -min_len_seed 5000, minimap2_options_cns=-t 8 -x ava-ont -k 17 -w 17 --minlen 1000 --maxhan1 5000. The NextDenovo2 assembly based on the HERRO-corrected data was performed in assemble mode without prior correction of the reads in NextDenovo2. BUSCO v5.7.1 (Simão *et al*., 2015; Manni *et al*., 2021) was run to assess the assembly completeness based on the eudicots_odb10 reference data set. Merqury v1.3 (Rhie *et al*., 2020) was run to assess the assembly correctness based on k-mer comparison between reads and the assembly.

### Annotation of the *Centaurea cyanus* gene set

RNA-seq reads (Additional File A) were retrieved from the Sequence Read Archive (Leinonen *et al*., 2011; Katz *et al*., 2022) via fastq-dump (NCBI, 2020). The alignment of these reads against the genome sequence was performed with STAR v2.7.11b (Dobin *et al*., 2013; Dobin & Gingeras, 2015) with default mapping parameters. Samtools v1.13 (Li *et al*., 2009) was applied to sort the BAM file by chromosomal position of alignments. GeMoMa v1.9 (Keilwagen,2016;Keilwagen,2019) was supplied with these RNA-seq hints and the following data sets for the prediction of protein-encoding genes: *Cichorium intybus* (GCA_023525715.1), *Ambrosia artemisiifolia* (GCA_024762085.1), *Helianthus annuus* (GCF_002127325.2) (Badouin *et al*., 2017), and *Lactuca sativa* (GCF_002870075.4) (Reyes-Chin-Wo *et al*., 2017).

Genes of the flavonoid biosynthesis were annotated using KIPEs v3.2.6 (Pucker *et al*., 2020; Rempel *et al*., 2023). MYB and bHLH transcription factors controlling the anthocyanin biosynthesis were identified with the MYB_annotator (Pucker, 2022) and bHLH_annotator (Thoben & Pucker, 2023), respectively. Candidates for the WD40 protein TTG1 were identified based on orthology to previously characterized sequences. Initial candidates were identified by BLASTp (Altschul *et al*., 1990, 1997) and integrated into a global alignment with previously characterized TTG1 and TTG1-like sequences using MAFFT v7 (Katoh & Standley, 2013).

FastTree v2.10 (Price *et al*., 2010) was used to infer a phylogeny based on this global alignment. The tree was visualized in iTOL (Letunic & Bork, 2024) for manual inspection of the candidates. A general functional annotation of all predicted protein-encoding genes was generated with a customized Python script construct_anno.py based on orthology to *Arabidopsis thaliana* TAIR10/Araport11 sequences (Lamesch *et al*., 2012; Cheng *et al*., 2017) as previously described (Pucker *et al*., 2017).

### Gene expression analysis

Publicly available RNA-seq data sets of *Centaurea cyanus* were retrieved from the Sequence Read Archive via fastq-dump. Quantification of gene expression was performed with kallisto v0.44 (Bray *et al*., 2016) based on these FASTQ files and the predicted coding sequences of the *C. cyanus* annotation.

## Results & Discussion

### *Centaurea cyanus* genome sequence and annotation

Based on a total of 32.5 Gbp with an N50 read length of 29.5 kb and an average Phred score of 38, assemblies were generated with different tools. These assemblies were compared with respect to contiguity, completeness, and accuracy (**Table 1**). The representative *C. cyanus* genome sequence comprises 720 Mbp spread over 391 contigs with an N50 length of 16.35 Mbp. BUSCO reported a completeness of 98.8 % based on the eudicots_odb10 and Merqury indicated a per base accuracy of 71.75.

**Table 1:** Statistics of different assemblies generated for *C. cyanus*. The assembly size, the number of contigs (>50kb), the accuracy reported by Merqury, and the completeness reported by BUSCO based on embryophyta_odb10 are listed for each assembly.

| ID | Size | Contigs | N50 | Merqury | BUSCO |
| --- | --- | --- | --- | --- | --- |
| GHN (HAC.nextdenovo) | 605 Mbp | 508 | 3.73 Mbp | 39.54 | 95.9% |
| GSN (SUP.nextdenovo) | 589 Mbp | 523 | 3.62 Mbp | 39.35 | 94.7% |
| DNN (DORADO.nextdenovo) | 623 Mbp | 492 | 4.22 Mbp | 39.60 | 96.3% |
| DNS (DORADO.shasta) | 677 Mbp | 2566 | 0.46 Mbp | 59.75 | 95.2% |
| DHN (DORADO.HERRO.nextdenovo) | 644 Mbp | 443 | 6.46 Mbp | 37.86 | 97.7% |
| DHS (DORADO.HERRO.shasta) | 720 Mbp | 391 | 16.35 Mbp | 71.75 | 98.8% |

Basecalling modes differing in their accuracy were compared to assess their influence on the final assembly result. Interestingly, the assembly based on high accuracy basecalling was slightly superior to the assembly based on super high accuracy basecalling with guppy (**Table 1**). It seems plausible that the relatively small difference is a random effect, but it is also possible that a stricter filtering of reads as part of the super high accuracy basecalling resulted in a smaller dataset. Dorado basecalling was performed for comparison, but did only slightly improve the assembly quality. A correction of the reads with HERRO resulted in a substantially improved assembly quality and caused Shasta to outperform NextDenovo2. The contiguity and completeness of the cornflower genome sequence on the contig level exceeds those of other genome sequences released for closely related species in its family (*Centaurea solstitialis* (Reatini *et al*., 2024), *Carduus pycnocephalus* (GCA_042919555.1),*Carthamus tinctorius* (Dong *et al*., 2024)). Therefore, we propose that this *Centaurea cyanus* genome sequence should be considered as a reference for future phylogenetic studies on the genus. Further improvement of the genome sequence would be possible based on Hi-C or Pore-C data.

Prediction of protein-encoding genes resulted in a completeness that correlates with the quality of the assembled genome sequence (**Table 2**). The numbers of around 28,000 protein-encoding genes align well with the average gene number expected in a plant genome (Pucker & Brockington, 2018).

**Table 2:** Structural annotation of the *Centaurea cyanus* genome sequence.

| Assembly | Number of genes | Annotation tool | BUSCO |
| --- | --- | --- | --- |
| GHN | 27036 | GeMoMa | C:94.9%[S:72.6%,D:22.3%],<br>F:0.5%,M:4.6% |
| DHN | 28171 | GeMoMa | C:96.9%[S:72.6%,D:24.3%],<br>F:0.4%,M:2.7% |
| DHS | 28763 | GeMoMa | C:98.7%[S:73.9%,D:24.8%],<br>F:0.2%,M:1.1% |

### Flavonoid biosynthesis genes in *Centaurea cyanus*

The predicted protein-encoding genes enabled the identification of the structural genes and transcription factors associated with the flavonoid biosynthesis (**Fig. 2**).

**Fig. 2:**
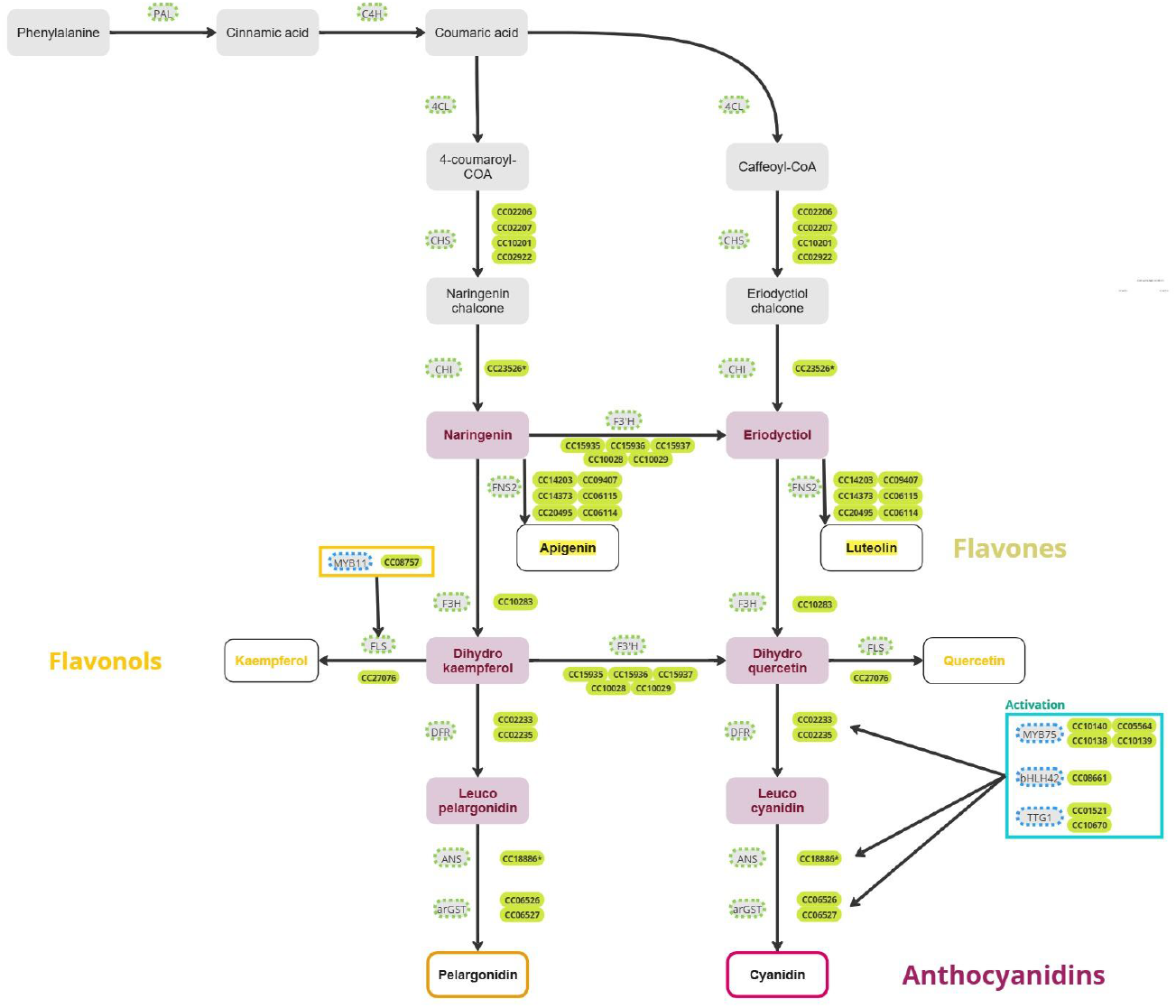
Simplified illustration of the flavonoid biosynthesis including candidate genes in *C. cyanus*. The formation of cyanidins and flavones is indicated as these are required as components of the protocyanin that is responsible for the striking blue color of cornflowers. Gene IDs marked with an asterisk have less than 100% conserved residues in their KIPEs results, but represent the best candidate for the respective step.

No *LAR* or *ANR* candidate genes were detected. Given the generally high assembly completeness and detection of many other genes in the flavonoid biosynthesis this suggests that *C. cyanus* does not have the canonical proanthocyanidin (PA) biosynthesis. To further investigate the PA biosynthesis genes, transcriptomic data from typically PA-rich plant parts such as seeds would be needed. Additionally, no F3 ‘5’H was found which is usually required to form delphinidin and produce blue flower colour. This supports the hypothesis that the blue colour is only an effect of the protocyanin. Genes encoding important enzymes such as the flavone synthase (FNS II) for the production of flavones (like apigenin) and the genes relevant for the accumulation of cyanidin (*DFR, ANS*, and *arGST*) have been identified (**Fig. 2**). Furthermore, genes encoding the components of the anthocyanin biosynthesis activating MBW complex (MYB75/PAP1,bHLH42/ TT8, and TTG1) (Gonzalez *et al*., 2008) were found. Although these genes could activate the pelargonidin pathway, it is highly unlikely that substantial amounts of pelargonidin are produced, because it could not be integrated into the protocyanin complex as it is missing a hydroxylation at the 3’ position on the B-ring (Schwinn *et al*., 2014). F3 ‘5’H is not present, as the cornflower uses protocyanin as its colour source, this is plausible as the activity of F3’5’H would take resources away. Comparison against a previously described *F3’H* transcript sequence derived from RNA-seq (Deng *et al*., 2019), revealed the corresponding gene in the annotation of our genome sequence. It appears that the gene in our assembly is still functional, which aligns with the observed pigments. As the polypeptide sequence derived from *CC15936* in our genome sequence only deviates by two amino acid residues from the F3’H candidate sequences reported in the previous transcriptome assembly, it is likely that this is the gene reported as crucial for the blue pigmentation by Deng et al., 2019. However, our assembly suggests that there are two copies of this gene located in tandem (*CC15935* and *CC15936*).

No perfect CHI candidate with 100% of the amino acid residues considered essential has been detected, but due to the existence of downstream genes and flavonoids it can be assumed that CHI functionality must exist in cornflower.

*DFR*, often considered as the first committed gene of the anthocyanin biosynthesis, can encode an enzyme that is specific for a substrate with a certain hydroxylation pattern of the B-ring (Choudhary & Pucker, 2024). An investigation of multiple cultivars and their respective DFR types showed high conservation among multiple cultivars (Deng *et al*.2019). We were able to support this finding as the DFR encoded in our genome sequence is not substrate specific according to previously reported determining amino acids (Choudhary & Pucker, 2024).

Furthermore, a gene encoding a flavonol synthase (FLS), a key gene for the flavonol biosynthesis, was identified. Studies have shown that the cornflower does produce flavonols (Litvinenko & Bubenchikova, 1988; Deng *et al*., 2019). As there is no indication that flavonols are present in the flower of *Centaurea cyanus* (blue cultivar), but flavonols were detected in other parts of the plants, we propose that there is an element regulating the *FLS* expression differentially throughout the plant. Further investigation through RNA-seq analysis of respective parts would be needed (Deng *et al*.2019). In total, the important genes for the formation of protocyanin are present.

Corresponding genes to previously reported anthocyanin-associated transcription factors were identified in our genome sequence: MYB6-1 (CC10140), MYB6-2 (CC10139), and bHLH1(CC08661) (Deng *et al*., 2020). The previously described MYB6-1 and MYB6-2 in cornflower (Deng *et al*., 2020) were classified as members of the PAP1 lineage thus suggesting that it is not a novel regulator (Additional File B). In addition, we identified CC10138 and CC05564 as members of the PAP1 lineage. The gene IDs suggest that CC10138, CC10139, and CC10140 are the product of a tandem gene duplication. By using primer sequences of the previously reported bHLH1 in cornflower (Deng *et al*., 2020), a gene that was classified as bHLH42 in a phylogenetic analysis conducted by the bHLH_annotator was identified. PAP1 MYBs are usually associated with TT8 (bHLH42) in the MBW complex (Gonzalez *et al*., 2008). This correlates with experiments in which both have been transformed together leading to substantial upregulation in anthocyanin biosynthesis and it is described that bHLH1 seems to be highly similar to TT8 (Deng *et al*., 2020).

## Declarations

## Competing interests

CD declares that Oxford Nanopore Technologies supplied the materials used for sequencing. RF and BP declare no conflict of interest.

## Data availability statement

All data sets underlying this study are publicly available. Sequencing data have been deposited at the European Nucleotide Archive (PRJEB76928). The assembled genome sequence, corresponding annotation, and analyzed gene expression data have been published via LeoPARD (https://leopard.tu-braunschweig.de/receive/dbbs_mods_00078333).

## Author contributions

CD and BP designed the project, conducted sequencing, and performed data analysis. RF conducted nanopore sequencing. All authors revised the manuscript and agreed to its submission.

## Acknowledgements

This work was supported by the BMBF-funded de.NBI Cloud within the German Network for Bioinformatics Infrastructure (de.NBI) (031A532B, 031A533A, 031A533B, 031A534A, 031A535A, 031A537A, 031A537B, 031A537C, 031A537D, 031A538A). We thank Vladislav Berg for excellent technical support. We thank all members of the research group Plant Biotechnology and Bioinformatics for discussion and support. We acknowledge support by the Open Access Publication Funds of Technische Universität Braunschweig.

## Additional information

Additional File A: RNA-seq data sets utilized as hints for the gene prediction process. Additional File B: Gene tree of the *Arabidopsis thaliana* and *Centaurea cyanus* MYBs.

## Notes

https://leopard.tu-braunschweig.de/receive/dbbs_mods_00078333

