## Supplementary figures and images for "Genome sequence of the blue flowering *Centaurea cyanus*"

### Additional File B

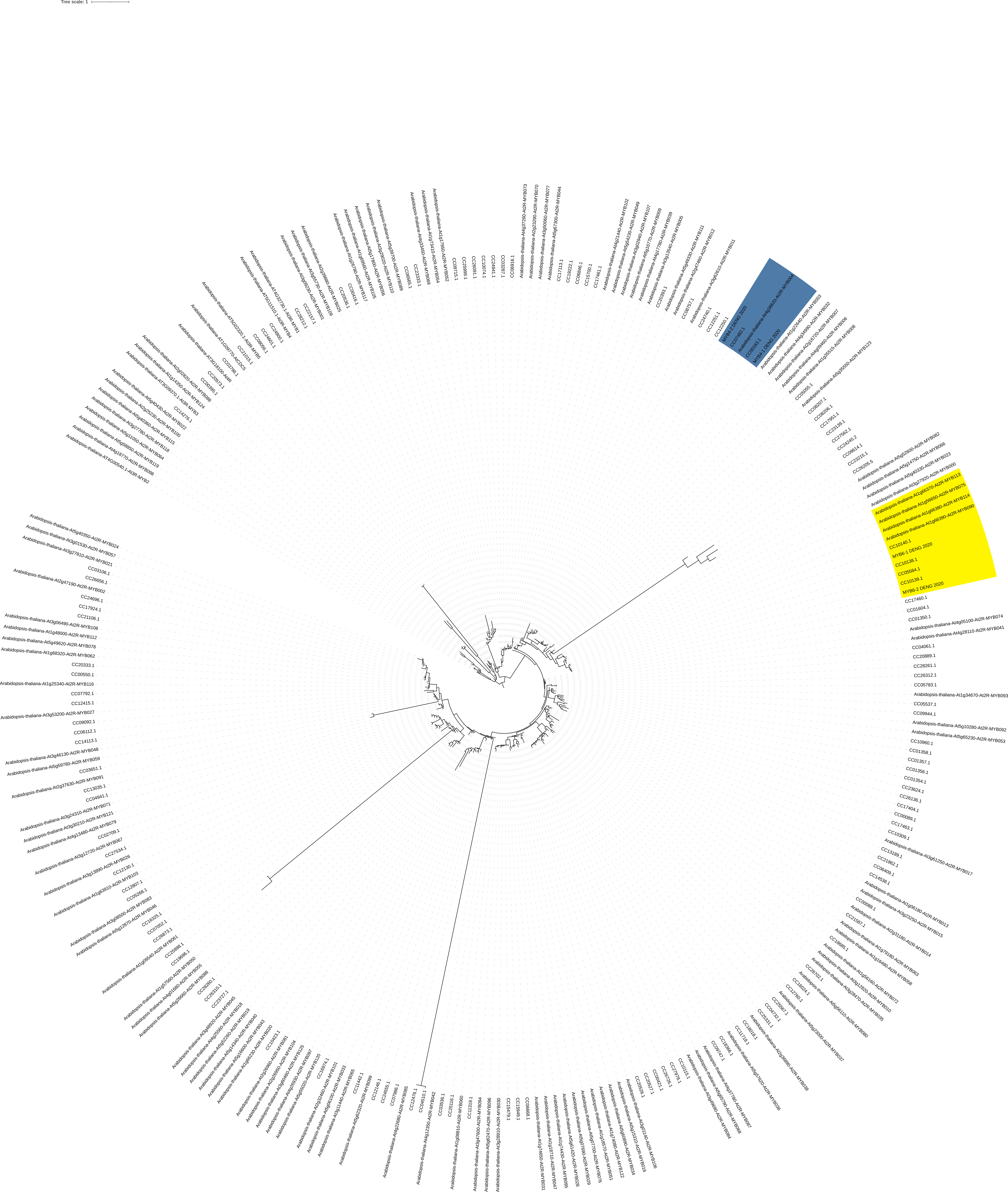
